# New components of the community based DNA-repair mechanism in Sulfolobales

**DOI:** 10.1101/2024.09.27.615169

**Authors:** Alejandra Recalde, Alexander Wagner, Shamphavi Sivabalasarma, Anastasiya Yurmashava, Nayeli Phycilia Fehr, Rebecca Thurm, Thuong Ngoc Le, Christin Köbler, Bianca Wassmer, Sonja-Verena Albers, Marleen van Wolferen

**Author notes:** **Correspondence** should be addressed to Marleen van Wolferen, or Sonja-Verena Albers.

## Abstract

After exposure to UV light, *Sulfolobus acidocaldarius* cells aggregate in a species-specific manner to exchange DNA and repair double-strand breaks via homologous recombination. The formation of cell-cell interactions is mediated by Ups pili. DNA exchange subsequently occurs through the Ced system, which imports DNA. To identify novel players in these processes, we investigated several genes upregulated after UV exposure by creating in-frame deletion mutants and performing cell aggregation and DNA exchange assays. This led to the identification of two novel components involved in the Ups and Ced systems: UpsC, a minor pilin of the Ups pili, and CedD, a VirD4-like ATPase essential for DNA import. Altogether, these findings provide new insights into the fascinating DNA damage response of Sulfolobales.

## Introduction

Ultraviolet (UV) light is one of the primary causes of DNA damage on Earth. Microorganisms, including archaea, have evolved various strategies to cope with this stress. These strategies include diverse DNA repair mechanisms (White & Allers, 2018), as well as protective measures such as biofilm formation (Lapaglia & Hartzell, 1997) and the production of photoprotective substances like carotenoids (Jones & Baxter, 2017). Other strategies include and DNA exchange (Grogan, 1998; van Wolferen et al., 2013a).

Upon DNA-damage caused by UV-light, species from the order of Sulfolobales aggregate using the Ups- system (UV inducible pili system). These type IV pili allow the cells to create cellular contacts (Fröls et al., 2008), and subsequently, they exchange DNA using the Crenarchaeal system for exchange of DNA (Ced) (Ajon et al., 2011; van Wolferen et al., 2016). Ups-pili are type IV pili encoded by the *ups-*operon that are present in all Sulfolobales (van Wolferen et al., 2013b). The *ups-*operon harbors: *upsE*, which encodes the assembly ATPase of the pilus; *upsF*, coding for the integral membrane protein of the pilus assembly machinery; *upsA* and *upsB*, both encoding pilin subunits that are processed by peptidase PibD before assembly into the pilus; and *upsX*, which codes for a protein of unknown function (Fröls et al., 2008; van Wolferen et al., 2013b). Deletion of any of the *ups-*genes, except for *upsX*, impairs cellular aggregation. Deletion of *ups*X, however, results in reduced DNA exchange (van Wolferen et al., 2013b).

Cellular aggregation of Sulfolobales was shown to be species-specific. This specificity is achieved by the interaction of the Ups-pili with the glycosylated S-layer of neighboring cells(van Wolferen et al., 2020). For this, pilin subunit UpsA is suggested to have a unique binding pocket that binds to the *N*-glycans that are specific for each species on the S-layer surface (van Wolferen et al., 2020).

In several bacteria, type IV pili have been shown to participate in the uptake of DNA(Filloux, 2010; Piepenbrink, 2019). Sulfolobales however, do not take up DNA from the environment, nor do they use the Ups-pili for DNA transfer(Ajon et al., 2011), rather they only exchange DNA between cells upon close cell-cell contact (Ajon et al., 2011; Fröls et al., 2008; Grogan, 1996). Once mating partners have been established using the Ups-pili, Sulfolobales species use the Ced system for DNA exchange(van Wolferen et al., 2016). Even though both systems are required for efficient DNA transfer, they form different complexes and do not need to be expressed in the same cell.

So far, four proteins of the Ced system have been described (van Wolferen et al., 2016): CedA, a protein containing 6-7 transmembrane domains, which is believed to form a channel similar to VirB6 in type IV secretion systems (T4SSs); CedB, a VirB4/HerA-like AAA+ ATPase that presumably powers DNA translocation across the membrane; and CedA1 and CedA2, both of which harbor two predicted transmembrane domains (van Wolferen et al., 2016). It has recently been described that CedA1 in *Aeropyrum pernix* (order Desulforococcales) and structural homolog TedC in *Pyrobaculum califontis* (order Thermoproteales), form a pili structure similar to VirB2 pilin in bacterial conjugation systems (Beltran et al., 2023). These findings thereby extend the Ced-system to the Thermoproteales where it is called Thermoproteales exchange of DNA (Ted) system (Beltran et al., 2023). In Sulfolobales however, no Ced- pili have so far been observed. The Ced and Ted system are thought to share a common ancestor with bacterial T4SSs conjugation systems (Beltran et al., 2023). Interestingly, the Ced system functions in the uptake of DNA rather than export as seen in bacterial conjugation systems (van Wolferen et al., 2016).

Several transcriptomic and proteomic studies regarding the UV-stress response and DNA damage in Sulfolobales have been performed (Feng et al., 2018; Fröls et al., 2007; Götz et al., 2007; Huang et al., 2020; Schult et al., 2018). Recently, a machine learning study combined transcriptomic datasets from *S. acidocaldarius* and identified genes regulated in a similar manner across different studies, grouping them into so-called iModulons (Chauhan et al., 2021). One of these iModulons was the TFB3-UV group, clustering genes that are upregulated upon UV stress dependent on general transcription factor TFB3. This transcriptional activator is expressed after dsDNA damage, triggering the expression of genes from the *ups* and *ced* operons (Feng et al., 2018; Le et al., 2017; Schult et al., 2018). Next to the *ups* and *ced* genes, several other non-characterized genes are part of this iModulon, including *saci_0667, saci_1302, saci_0951, saci_1225, saci_1270* and *saci_RS06010*.

Here, we investigated all previously unstudied genes of the UV-TFB3 iModulon and created in-frame deletion mutants to explore their involvement in the UV-light response. Using aggregation- and DNA transfer assays, two novel components of previously studied systems were found: *saci_1302 (upsC)*, encoding an additional pilin subunit of the Ups pili, which is necessary for cell aggregation; and *saci_0667 (cedD*), an additional component of the Ced-system essential for DNA transfer.

## Results

### Bioinformatics analysis

Among the genes that show TFB3-dependent regulation upon UV-stress, there are the *ups* genes encoding the Ups-pili and the *ced* genes encoding the Ced-system for DNA transfer (Table 1). To study all other genes that are part of the TFB3-UV iModulon, we performed bioinformatics analyses and created deletion mutants to analyze their putative roles.

**Table 1.**
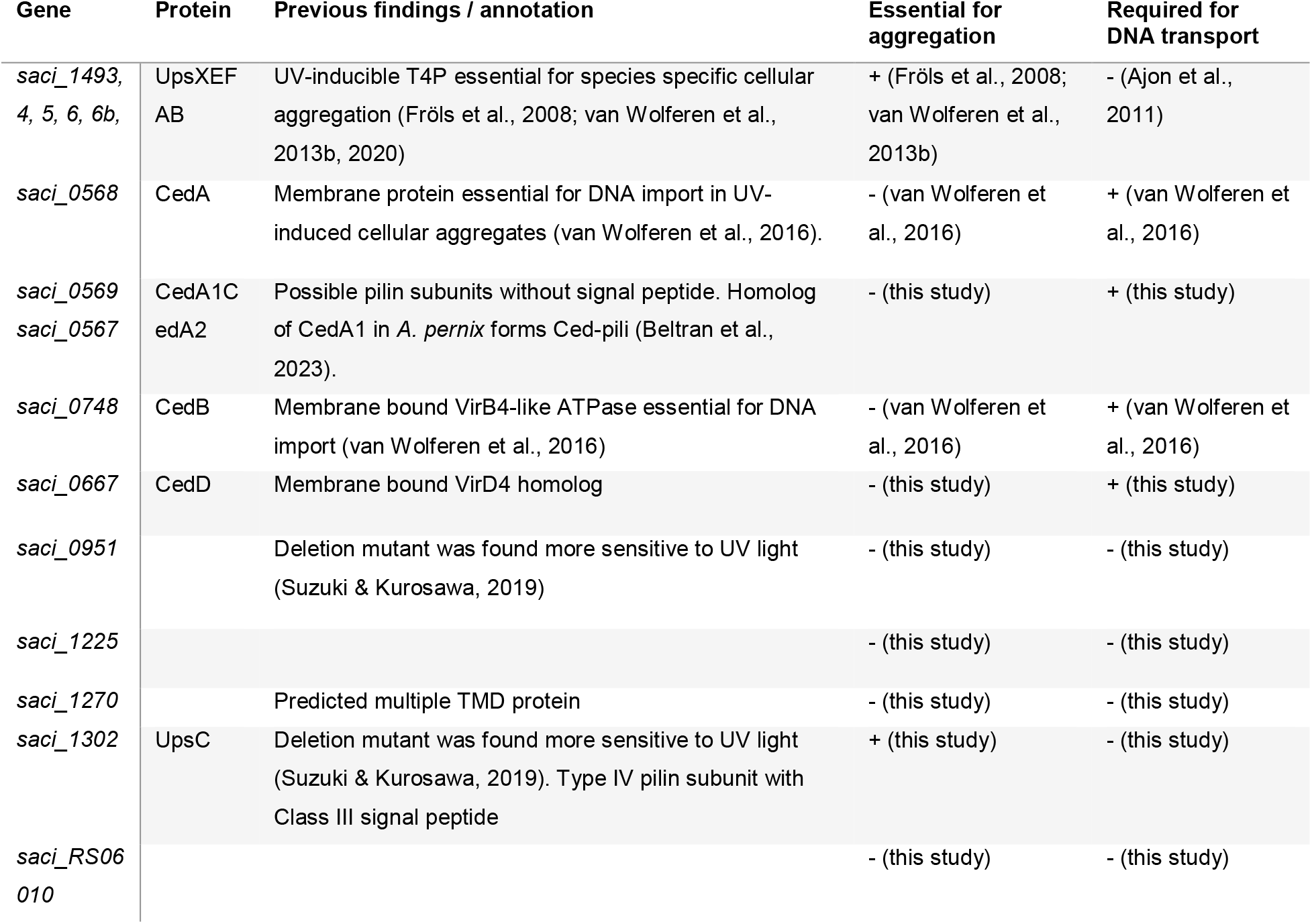
Upregulated genes in *S. acidocaldarius* upon UV stress. A summary of previous findings and results from this study, indicating whether these genes are essential for cellular aggregation and/or DNA transport. Homologs from other species are listed in Figure S1.

**Table 2.** Genotypes of recombinants from UV-treated *ΔcedD* and *ΔupsE* mixtures. Colony PCR was performed on the colonies resulting from the UV*UV mating experiments shown in Figure 3.

| Mixture | Gene | Colonies |  | Gene present in |
| --- | --- | --- | --- | --- |
|  |  | + | Δ |  |
| *ΔcedD\*ΔupsE* | *cedD* | 35 | 0 | 100% |
|  | *upsE* | 9 | 26 | 26% |

Similar to the *ups*-genes, *saci_0951, saci_1225, saci_1270 saci_1302* and *saci_RS06010* are exclusively found in Sulfolobales. *Saci_0667* can additionally be found in the Desulfurococcales and Acidilobales, as are the *ced* genes (van Wolferen et al., 2016) (Table S1).

Genes *saci_0951, saci_1225* and *saci_RS06010* all encode small proteins that are predicted to bind DNA, they might therefore play a role in transcriptional regulation upon UV-induction. Saci_1270 is a multi-pass transmembrane protein conserved across all Sulfolobales. It has been hypothesized to play a role in DNA export by working in conjunction with the Ced system (Chauhan et al., 2021).

AlphaFold models of CedA1 and CedA2 (encoded next to *cedA*, Figure 1A) show a structure similar to VirB3 (Figure 1B), making it very likely that they are pilin subunits, as shown for CedA1 in *A. pernix* and homolog TedC in *P. calidifontis* (Beltran et al., 2023). Interestingly, in the Sulfolobales and Desulforococcales, CedA1 and CedA2 do not harbor any signal peptides, whereas their homolog TedC in the Thermoproteales contains a Sec signal peptide.

**Figure 1.**
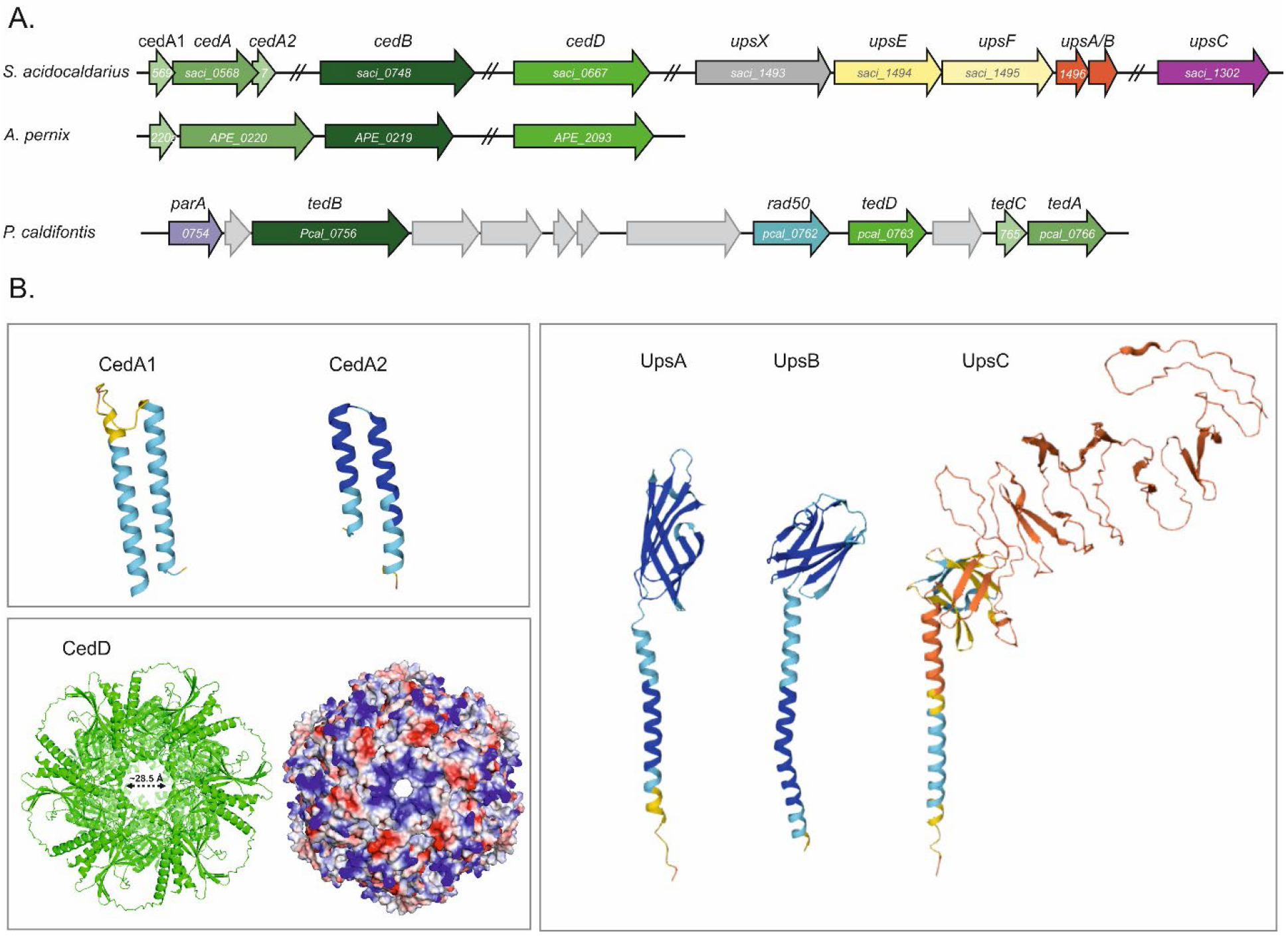
(A) Genetic loci of the *ced* and *ups* genes of *S. acidocaldarius* and *A. pernix*, as well as the *ted* genes of *P*.*caldifontis*. (B) AlphaFold predictions of *S. acodocaldarius* proteins, including: the putative pilin subunits CedA1 and CedA2; a hexamer of CedD; as well as pilin subunits UpsA, UpsB and UpsC from *S. acidocaldarius*. The hexamer of CedD was modelled using the AlphaFold server and has a predicted pore diameter of around 28 Å (left). Positive and negative charges are represented in blue and red, respectively, revealing a positively charged inner pore (right).

Similar to *cedB, saci*_*0667* (now called *cedD*) encodes a membrane-bound ATPase belonging to the HerA- FtsK superfamily. Based on homology we previously proposed that CedB functions similarly to VirB4 in T4SSs, energizing the biogenesis of the Ced system and facilitating DNA translocation (van Wolferen et al., 2016). CedD might in turn play a role analogous to VirD4 as a coupling protein, as both ATPases are essential in bacterial conjugation systems.(Costa et al., 2024). Supporting this hypothesis, KEGG and SMART databases predict a coupling protein DNA-binding domain in CedD. Modelling CedD as a hexamer revealed an inner pore of around 28 Å with positive charges, theoretically allowing translocation of ssDNA as well as dsDNA (Figure 1B).

To search for homologs of CedD, we used EggNOG, KEGG, SyntTax and BLAST (Iyer et al., 2004). Similar to CedB, in some organisms CedD lacks a transmembrane domain at its N-terminus, a feature frequently observed in VirB4 homologs in Bacteria (van Wolferen et al., 2016) (Table S1). These ATPases might, therefore, bind to the membrane via other factors, possibly CedA, in a way similar to T4SS ATPases via VirB3 (Fronzes et al., 2009). In Thermoproteales, where the Ted pili were isolated (Beltran et al., 2023), we also identified homologs of CedD, which we named TedD following the Ted nomenclature. Interestingly, TedD is encoded within the same genetic locus as the other Ted proteins, supporting the hypothesis that it is part of the same system (Figure 1A, Table S1)

*Saci_1302* (now named *upsC*) encodes putative type IV pilin subunit (Sayers et al., 2022). It co-occurs with the *ups* genes within the Sulfolobales but is not located within the *ups* operon (Figure 1A, Table S1). In addition to the typical N-terminal class III signal peptide (Figure S1A), it harbors an unusually large C-terminal domain (protein size of 49 kDa compared to 18 kDa for UpsA). BLAST analysis does not provide any insights towards the function of this domain. Unlike the AlphaFold models of pilin subunits UpsA and UpsB, the C-terminal domain of UpsC has a very low model confidence (Figure 1B), making it also difficult to predict any putative functions for this domain based on its structure. Similar to other pilin subunits and surface proteins of *Sulfolobus* (Gaines et al., 2024; Meyer et al., 2015; Sofer et al., 2024), UpsC is heavily glycosylated and harbors 9 predicted glycosylation sites in its predicted globular domain.

### Pilin subunit UpsC is essential for cellular aggregation

Because In order to study the genes that are part of the TFB3-UV iModulon, deletion mutants of *cedD* (*saci_0667*), *ups*C (*saci_1302*), *saci_1225, saci_1270, saci_RS06010* and *saci_0951* were created using the pop-in/pop-out method (Wagner et al., 2012) (Table S2 and S3). In a previous study, a deletion mutant of *saci*_*0951* was found to be sensitive to UV radiation (Suzuki & Kurosawa, 2019). In our hands however, we were unable to delete this gene entirely, likely because a complete deletion would disrupt the promoter of the neighboring gene, *saci_0952*, which is essential in *S. islandicus* (Zhang et al., 2018). Therefore, we created a partial deletion mutant that retains the first 81 base pairs of *saci_0951*. Additionally, we also created deletion mutants of *cedA1* (*saci_0569*) and *cedA2* (*saci_0567*), encoding the putative Ced pilin subunits, analogous to TedC in the Ted-pili of *A. pernix* (Beltran et al., 2023).

To test if any of the deleted genes are involved in the formation of cell-cell contacts, we performed aggregation assays on the derived mutants. All deletion strains showed normal cellular aggregation after UV-light exposure, except for Δ*upsC*, which did not aggregate (Figure 2AB and Figure S2), consistent with previous findings for other *ups* deletion strains (Fröls et al., 2008; van Wolferen et al., 2013b). Aggregation could be restored when complementing Δ*upsC* with an HA tagged or non-tagged version of UpsC expressed from a plasmid (Figure 2A). To determine whether the deletion of *upsC* also affects Ups-pili formation, we deleted *upsC* in a strain lacking Aap pili and archaella (MW1351) (Table S2). Electron microscopy of negatively stained cells revealed that pili were still formed, though in reduced amounts, indicating that UpsC is not essential for pili formation (Figure 2C).

**Figure 2:**
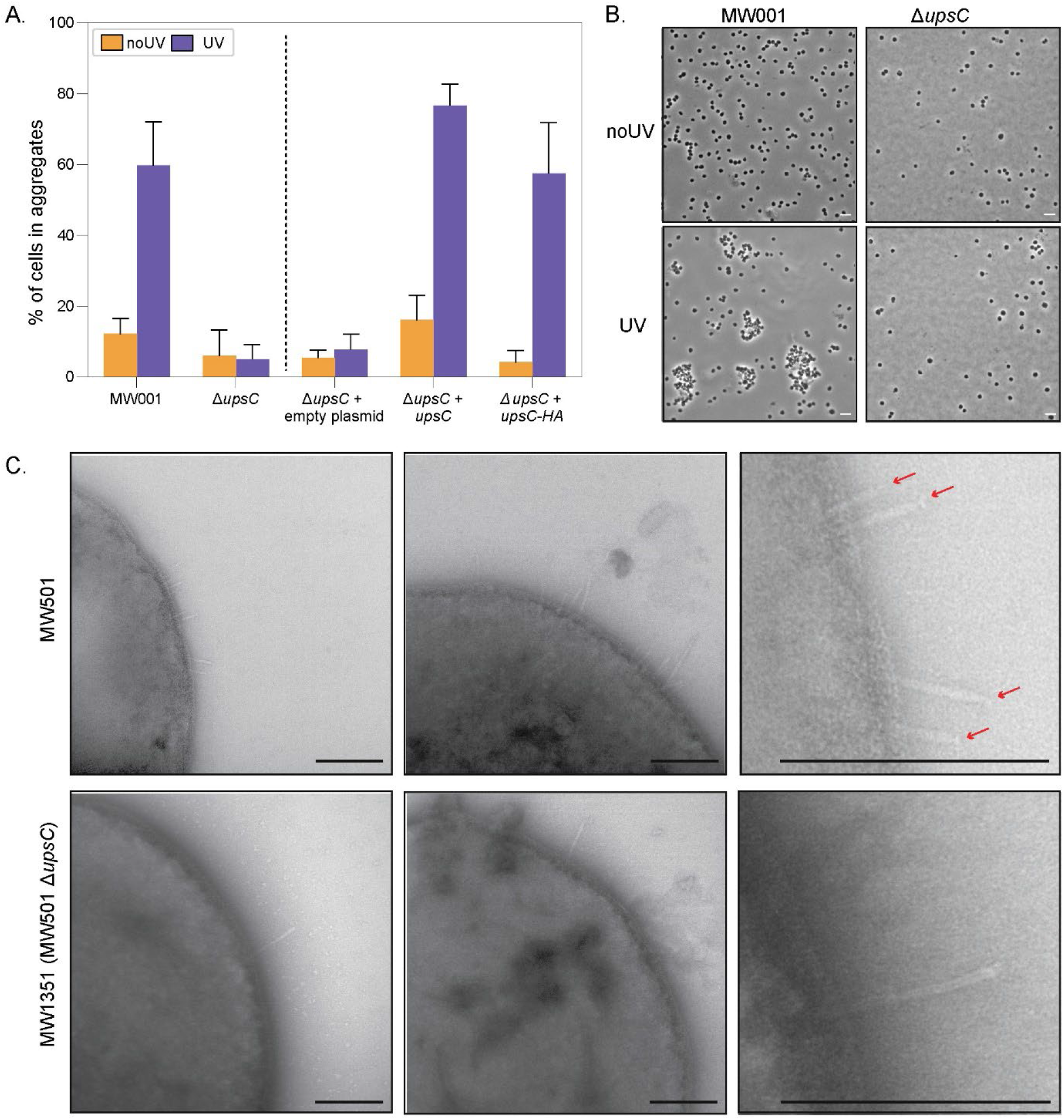
UV induced cellular aggregation *S. acidocaldarius* MW001, Δ*upsC* and complementation strains expression *upsC* and *upsC-HA* from a plasmid. (A) Percentages of aggregated cells, after UV light exposure and the non-exposed control. (B) Representative phase contrast images of MW001 and Δ*ups*C with and without UV treatment. Scale bars = 5 µm. (C) Electron microscopy images of MW501 (lacking Aap pili and archaella) and MW1351 (ΔupsC in MW501) strains. Ups pili are visible in both strains, showing that pilus formation is reduced but not abolished in the absence of UpsC. Red arrows point to a putative additional structure at the tip of the Ups-pili in MW501. Scale bars = 200 nm.

Based on its annotation and predicted structure, we suggest Saci_1302 to be a minor pilin of the Ups-pilus possibly involved in forming cell-cell interactions and named it UpsC. In line with this, we showed that the class III signal peptide of UpsC is processed by signal peptidase PibD in heterologous cleavage assays (Figure S1A and B). As expected, we moreover observed membrane localization, when separating membrane from cytosol from *S. acidocaldarius* cells expressing the HA tagged version of UpsC (Figure S1C). The major fraction of UpsC-HA runs higher than the expected size of 49 kDa which is probably due to the mentioned glycosylation of the pilin subunit

### CedD is a new component of the Ced-system

To test whether any of the other genes of interest are involved in the UV-light response, we performed DNA transfer assays. For this, *upsC (saci_1302), cedD (saci_0667), saci_0951, saci_1225, saci_1270, saci_RS06010, cedA1* (*saci_0569*) and *cedA2* (*saci_0567*) deletion mutants (Table S1) were generated in background strains MW001 and JDS22. MW001 and JDS22 are two different uracil auxotrophic strains with non-overlapping deletions in *pyrE*: MW001 (311 bp deletion; nt 91–412 in *pyrE*) (Wagner et al., 2012) and JDS22 (22 bp deletion; nt 16–38 in *pyrE*) (Grogan & Hansen, 2003). When the two background strains (MW001 and JDS22) are mixed together, the exchange of chromosomal DNA via homologous recombination enables the restoration of *pyrE*, resulting in prototrophic colonies that can grow without uracil. As previously observed (Ajon et al., 2011), inducing one of the two strains with UV light (UV*UV) led to an increase in the number of recombinants, and inducing both strains resulted in an even higher number of recombinants, which was set to 100% (Figure 3).

**Figure 3.**
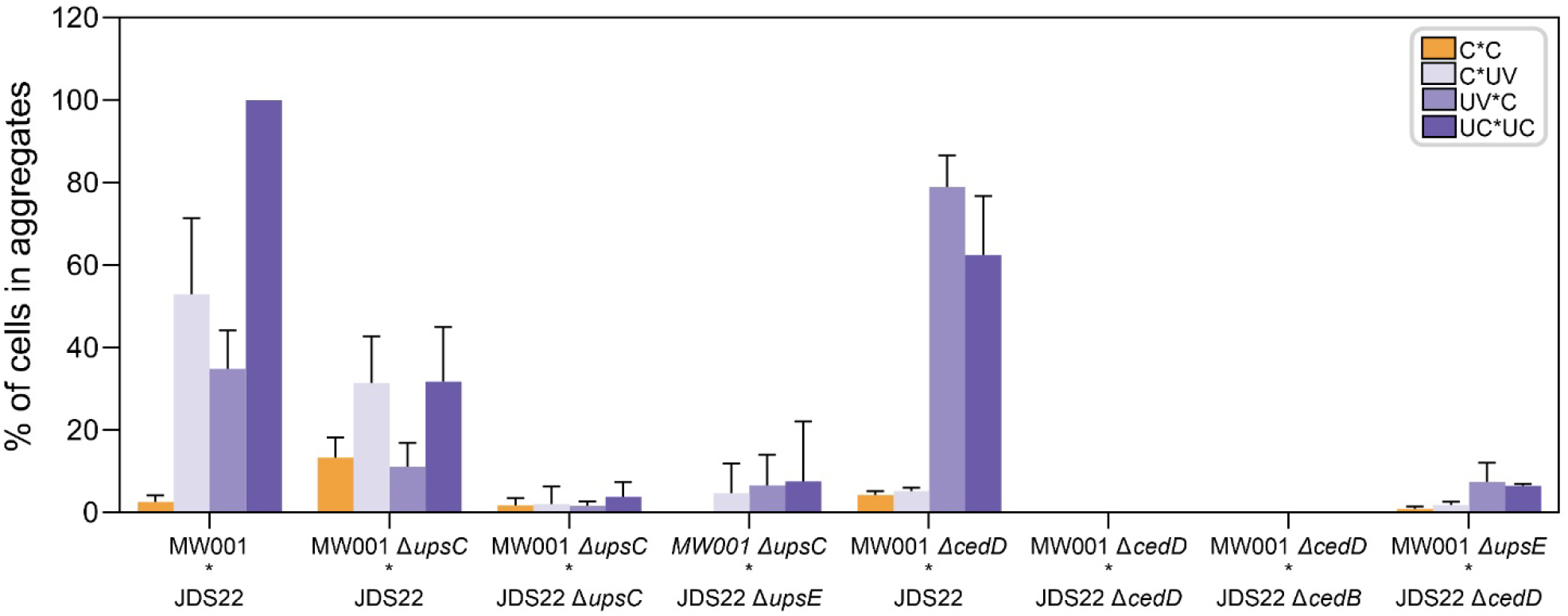
DNA exchange assays using *upsC, cedD, upsE* and *cedB* deletion mutants. Two different mutants (either in JDS22 or MW001 backgrounds), exposed to UV light (UV) or left untreated (C), were mixed in various combinations and plated on selective media. Both background strains carried mutations in the *pyrE* gene, which is involved in uracil biosynthesis. The mutations were located at different genomic positions, allowing recombination between the strains to restore the wild-type *pyrE* phenotype. The bars represent the average of at least three independent mating experiments, with each experiment normalized to the JDS22 (UV) × MW001 (UV) condition, set at 100%.

None of the deletion mutants *Δsaci_1225, Δsaci_1270, Δsaci_RS06010*, and *Δsaci_0951* showed a reduced recombination rate compared to the background strains, suggesting that these genes are not involved in DNA transfer. Therefore, the previously hypothesized role of *Saci_1270* in DNA export could not be confirmed (Figure S3).

Similar to what we observed in DNA transfer assays with other *ups* deletion mutants (Ajon et al., 2011) and in line with the aggregation assays, little to no DNA transfer was observed upon mixing two Δ*upsC* strains (Figure 3). When mixing *upsC* with *upsE*, similar results could be observed, suggesting UpsC to be part of the Ups system. Moreover, when mixing Δ*upsC* with background strain JDS22, an increase in recombinants was only observed when inducing JDS22 with UV light (Figure 3).

As previously described for the *cedA* and *cedB* deletion mutants (van Wolferen et al., 2016), a mixture of two Δ*cedD* strains did not result in any colony formation (Figure 3), indicating that CedD is essential for DNA transport. Importantly, when mixing a Δ*cedD* with a Δ*cedB* strain, no colonies were formed either, suggesting that both proteins operate within the same machinery. In contrast, in a mixture of Δ*cedD*\*Δ*upsE*, the Δ*upsE* cells are still able to transfer DNA, while the Δc*edD* cells still express Ups pili, facilitating the formation of mating partners. Consequently, a small number of recombinants can still be observed in this mixture (Figure 3).

To confirm that CedD is also involved in DNA import, we conducted colony PCR on recombinants resulting from the mixture of Δ*cedD* and Δ*upsE* (Figure 3). We found that the *cedD* gene was present in 100% of the recombinants, while the majority (74%) did not contain *upsE* (Table 1). This indicates that only *cedD*-positive cells are capable of importing DNA. The 26% of cells that carried both genes probably resulted from a secondary DNA transfer or recombination event, as previously observed in the *ΔcedA*\**ΔupsE* mixtures (van Wolferen et al., 2016).

For completeness, we also conducted DNA exchange experiments on the deletion mutants of *cedA1* and *cedA2* (Figure S3). CedA1 was shown to form Ced-pili in *Aeropyrum pernix* (Beltran et al., 2023). The deletion of these genes resulted in no colony formation in the mixes *ΔcedA1*ΔcedA1* and *ΔcedA2*ΔcedA2*, making it highly likely that CedA1 and CedA2 also function as pilin subunits in Sulfolobales. Given the fact that we have never observed any Ced-pili, and because cellular interactions are achieved by the Ups-pili, it is likely that they rather form pseudo-Ced pili in these species.

## Discussion

Several microorganisms take up DNA from the environment to maintain genome integrity upon DNA damage (Ajon et al., 2011; Claverys et al., 2006). Unlike bacteria, Sulfolobales cannot take up free DNA (Grogan, 1996) ; instead, DNA uptake occurs via the Ced system (van Wolferen et al., 2016), which can only occur upon close cell-cell contact mediated by the Ups pili (Ajon et al., 2011).

The response of Sulfolobales to UV stress has been extensively studied. In this work, we aimed to identify additional genes that are highly upregulated upon UV stress and actively participate in cell aggregation, DNA exchange, or related processes. By creating deletion mutants of target genes in different background strains, we identified two new components of the Ups and Ced systems.

During this study, we identified an additional type IV pilin subunit, UpsC, belonging to the Ups system. UpsC is also essential for cellular aggregation and is conserved among all species that possess a functional Ups system. Notably, UpsC is not encoded within the *ups* operon (van Wolferen et al., 2013b). The previously observed decreased survival rates in *upsC* (*saci_1302*) mutants (Suzuki & Kurosawa, 2019), are likely due to impaired cell aggregation, leading to significantly reduced DNA exchange rates. UpsC, with 49 kDa, is more than three times the size of the pilin subunits UpsA and UpsB, which are 15 and 14 kDa, respectively. Both UpsA and UpsB are thought to be the major pilin subunits of the Ups pili (van Wolferen et al., 2013b), with UpsA also shown to play a role in species-specific binding of glycosylated S-layer proteins from cells of the same species (van Wolferen et al., 2020). In contrast, UpsC is probably a minor pilin subunit. It may play a role in the initiation of pilin formation at the base of the complex or function in initial adherence to other cells as part of the tip complex as is the case for other minor pilin (Jacobsen et al., 2020).

Interestingly, GspK from type II secretion systems (analogous to T4P) is a similarly large minor pilin (Korotkov & Hol, 2008). GspK has a large insertion in its globular domain, which was proposed to ensure that it is incorporated only at the tip of the pseudopilus. This increased size might play a role in shielding the cavity at the tip (Korotkov & Hol, 2008). In several EM images from this and previous studies, a different density appears to be visible on the Ups pili (Figure 2C, red arrows). Therefore, UpsC could play a role similar to GspK, potentially positioning at the tip of the pilus. Here, UpsC might also be involved in the initial recognition of other cells. To validate this hypothesis, further analysis of additional EM images will be essential.

Because we now know that archaeal T4P can retract (Charles-Orszag et al., 2024), Ups pili might also be able to do so. After the initial interaction between cells, the retraction of the pili could bring the cells into closer proximity allowing the transfer of DNA between cells via the Ced system.

We additionally identified CedD as a novel component of the Ced system. Similar to other Ced components, CedD is essential for DNA transfer and functions in DNA import. CedD is a membrane-bound ATPase belonging to the FtsK-HerA superfamily. Orthologs of this protein are found in organisms with either a Ced or a Ted system; in the latter, we named it TedD. The fact that it is located near the other *ted* genes, further supports the likelihood that they function together. Other genes within this locus might encode additional proteins involved in the Ted system and will be of particular interest for future studies. These include genes encoding DNA processing proteins such as ParA or Rad50, indicating that these proteins may also be involved in DNA processing or transfer, as proposed before (Beltran et al., 2023).

In bacterial T4SSs, ATPases such as VirD4 and VirB4 work together to coordinate DNA transfer through the inner membrane (Christie, 2016). In the Ced system, CedB likely energizes both the assembly of the Ced system and possibly the internalization of DNA (van Wolferen et al., 2016). We hypothesize that CedD functions as a coupling protein analogous to VirD4; however, instead of preparing DNA for export, it would guide incoming DNA during uptake, forming a critical link between the DNA and the Ced system. Additionally it could be involved in gating by opening the translocation pore.

It is likely that other currently unstudied proteins play a role in the Ced system. While T4SSs have primarily been studied in gram-negative bacteria, components that interact with or pass through the outer membrane are likely absent in archaea. Instead, components binding the S-layer may be present and will be of interest in future studies to elucidate the entire Ced system. Whether there is an export component acting in the other cell of the mating pair is still unclear. The additional highly upregulated genes analyzed in this study were not found to be essential for either mating pair formation or DNA transfer, suggesting they are probably not part of the Ced or Ups systems.

## Model and Conclusion

Based on the findings described in this paper and previous studies, we propose the following model (Figure 4). Upon UV treatment, Sulfolobales form mating pairs via the Ups pili. The minor pilin UpsC may form a tip complex, potentially involved in initiating mating pair formation, major pilin UpsA subsequently helps in forming tighter interactions through interaction with the S-layer (van Wolferen et al., 2020). Other species, such as the Thermoproteales, lacking the Ups system may form mating pairs directly using Ced (or Ted) pili (Beltran et al., 2023). Key elements of the Ced system—CedA (a VirB6 homolog), CedB (a VirB4 homolog), and CedA1 (a VirB2 homolog)—suggest a machinery reminiscent of bacterial Type IV Secretion systems, particularly in the formation of a membrane-spanning channel and pilus structure. Genomic DNA transfer is likely initiated at sites of DNA damage. While it is still unclear whether single or double stranded DNA is transferred and a dedicated relaxase enzyme has not been identified, DNA repair proteins such as HerA and NurA may prepare the DNA for transfer. Unlike bacterial conjugation systems, which export DNA, the Ced system facilitates direct DNA import from one cell to another. The core components of the DNA translocation machinery include CedA, which forms the translocation channel, and CedA1 (and possibly CedA2), which forms a (pseudo)pilus that spans the membrane and contacts the recipient cell. Energy for the assembly is provided by the ATPase CedB. VirB4 homolog CedD might then open the translocation pore and pull the DNA through the Ced system. Incoming DNA would then undergo repair via homologous recombination. However, the exact pathway of DNA processing remains unclear, as does the mechanism at the donor site of DNA transfer. Several T4SS proteins found in bacteria have not been so far found in archaea, suggesting that other unknown players are likely involved and yet to be identified. The distribution of the Ced and Ted systems suggests a common DNA transfer and repair mechanism among the Sulfolobales, Desulfurococcales, and Thermoproteales, indicating that it might be a common trait in thermoproteota.

**Figure 4.**
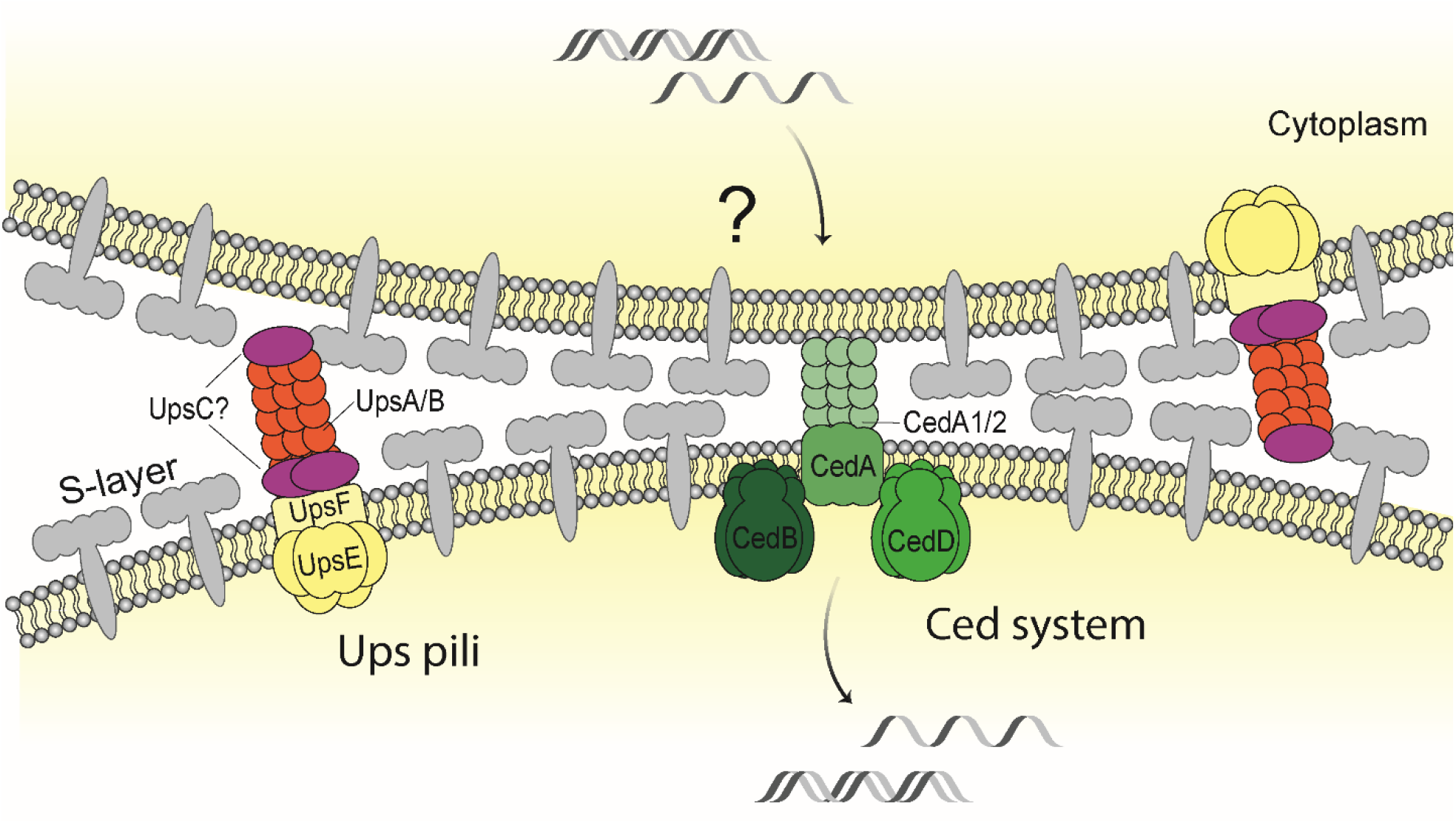
DN Current model of DNA transport in Sulfolobales via the Ced system. *Sulfolobus* cells aggregate after UV-stress, a process facilitated by the Ups pili system. UpsC, a minor pilin subunit, may function either at the base or tip of the Ups-pilus, with a potential role in initial cell adherence if positioned at the tip. The Ced system enables the import of DNA from neighboring cells, which is mediated by CedA, a polytopic membrane protein. CedA1 and CedA2, likely functioning as pseudopilins, may form a channel through which DNA is transferred between cells. CedB, a membrane-bound VirB4-like ATPase, and CedD, a VirD4-like membrane-bound ATPase, collaborate to power the DNA transport process. CedB likely energizes the assembly of the DNA transport machinery, while CedD functions as a coupling protein, processing incoming DNA and guiding it through the channel formed by CedA, CedA1, and CedA2. CedD may also act as a gatekeeper, controlling the opening of the translocation channel. It remains unclear whether the Ced system transports single-stranded or double-stranded DNA. Once inside the recipient cell, the imported DNA is used for DNA repair via homologous recombination.

In this study we identified UpsC as an additional minor pilin subunit of the Ups pilus and CedD as a VirD4-like ATPase in the Ced system. We have thereby discovered two novel players involved in this fascinating community-based DNA repair mechanism found amongst archaea.

## Material and methods

### Bioinformatics analysis

Homology searches of our proteins of interest were done using BLAST (Altschul et al., 1990), SyntTax (Oberto, 2013) and EggNOG (Huerta-Cepas et al., 2019). Proteins were modeled and visualized using AlphaFold3 and the AlphaFold Server (Varadi et al., 2024) and PyMOL (Schrödinger, LLC, 2015). Transmembrane domains were predicted using DeepTMHMM (Hallgren et al., 2022). For prediction of DNA binding, we used DNABIND (Szilágyi & Skolnick, 2006). Class III signal peptide cleavage sites were predicted using FlaFind 1.2 (Szabó et al., 2007). N-glycosylation sites were predicted using the NetNglyc 1.0 server (Gupta & Brunak, 2002).

### Strains and culture conditions

*S. acidocaldarius* MW001, JDS22, and uracil auxotroph-derived mutants (Table 3) were grown in Brock media (pH 3.5) (Brock et al., 1972) supplemented with 0.2% (w/v) sucrose, 0.1% (w/v) N-Z amine (Sigma-Aldrich®, Merk KGaA, Burlington, MA, USA), and 10 μg/mL uracil aerobically at 75°C. Strains carrying plasmids were grown without uracil, and expression was induced with 0.2% (w/v) xylose. For solid media, first selection plates contained gelrite (1.2%), 20 mM MgCl_2_, and 6 mM CaCl_2_ in unadjusted Brock medium supplemented with 0.2% (w/v) dextrin and 0.1% (w/v) N-Z amine. Second selection plates included 100 μg/mL 5-FOA and 10 μg/mL uracil. Competent *E. coli* DH5α and ER1821 (NEB) were used for cloning and methylation of plasmid DNA, respectively, and were grown at 37°C in LB supplemented with appropriate antibiotics.

### Construction of in-frame markless deletion mutants in *S. acidocadarius*

To construct deletion strains of *S. acidocaldarius* MW001 and JDS22 (Grogan & Hansen, 2003; Wagner et al., 2012), plasmids were generated following the method described by (Wagner et al., 2012), using primers listed in Table S2. In brief, around 500 bp upstream and downstream regions from each gene were amplified and connected via overlap PCR. The fragment was subsequently cloned into pSVA431, resulting in the plasmids listed in Table S2. Plasmid transformation, initial selection via blue/white screening, secondary selection, and mutant screening were performed as previously described (Wagner et al., 2012). The deletion mutants were confirmed by sequencing.

### UV light exposure and aggregation Assays

*Sulfolobus* strains were grown in 50 mL until an OD_600_ of 0.2–0.4. The cultures were then split in half, with 25 mL treated with UV at 75 J/m^2^ in plastic petri dishes before being returned to Erlenmeyer flasks. The other half of the culture was kept as a control. Both cultures were incubated for 3 hours at 75°C and 110 rpm.

To assess aggregation of the cells, 3 μL of culture were spotted on agarose pads (Basal Brock, 2% agarose) and allowed to dry. A coverslip was then placed over the pads, and the samples were observed under a Zeiss Axio Observer Z1 phase contrast microscope equipped with a Plan-Apochromat 100x 1.40 Oil Ph3 M27 objective. Three images per condition and strain were taken from each of three independent experiments. The percentage of cells in aggregates was calculated by counting all the cells using ImageJ software. (NIH, Bethesda, MD) (Schindelin et al., 2012).

### Cleavage assays

For the cleavage assays, a truncated version of the N-terminal portion of UpsC was heterologously expressed in *E. coli*. Controls included plasmids pSVA906 and pSVA914, containing *aapA*-HA and *aapA*-HA + *pibD*-His, respectively (Henche et al., 2014).

Plasmids pSVA13169 and pSVA13197 (Table S2) were created for truncated UpsC expression with or without signal peptidase PibD, respectively. Plasmid pSVA13169 was constructed using in vivo assembly, amplifying pSVA914 without the *aapA* gene with primers 12094-5, followed by DpnI treatment to remove the template. The N-terminal region of *saci_1302* was amplified from *S. acidocaldarius* genomic DNA using primers 12096-7. The purified linear PCR products were then transformed into *E. coli* DH5α cells. For pSVA13197, primers 13235-6 were designed to linearize pSVA13169, leaving out the *pibD*-His gene. The linear fragment was phosphorylated, ligated, and transformed into *E. coli* DH5α. Plasmids were purified from individual colonies and sequenced for verification.

Plasmids were transformed into *E. coli* BL21 (DE3) RIL and grown overnight at 37°C in LB medium with ampicillin and 1% glucose. An aliquot was transferred to fresh LB-ampicillin media, and when the OD reached 0.2, Saci_1302-HA or AapA-HA production was induced with 0.2% L-arabinose. After 2 hours, PibD-His production was additionally induced with 0.25 mM IPTG. One culture was left uninduced as a control. After 2 more hours, cells were collected by centrifugation.

The cells were resuspended in 4 mL HEPES buffer (50 mM HEPES, pH 7.5, 500 mM NaCl, 10% glycerol, 1X protease inhibitor cocktail) and passed through a French pressure cell at 800 lb/in^2^. Unbroken cells were removed by centrifugation, and the membrane fraction was collected by ultracentrifugation (250,000 × g, 45 min at 4°C) and resuspended in 100 μL HEPES buffer.

Samples were mixed with protein loading dye and boiled at 95°C for 5 min. Proteins were separated on two 15% SDS-PAGE gels, stained with Coomassie, or blotted onto a PVDF membrane. Blocking was done overnight at 4°C with a 0.1% I-Block solution in PBS-T. The primary α-HA antibody (rabbit) (Sigma-Aldrich) was used at a 1:10,000 dilution for 4 hours, followed by washing. The secondary anti-rabbit HRP antibody (1:10,000 dilution, Sigma-Aldrich) was applied and incubated for 3 hours. Chemiluminescence was initiated with HRP substrate (Clarity Max Western ECL Substrate, BioRad), and signals were imaged using the iBright FL1500 system (Invitrogen).

### Localization of UpsC

For localizing UpsC-HA, *S. acidocaldarius* MW1351 was transformed with plasmid pSVA13161, Cultures with an OD of 0.2 were induced with 0.2% (w/v) xylose for 1 hour, then treated with UV-light as described above. Control cells contained the empty plasmid pSVAaraFX-HA. After 3 hours, 1.5 mL of cells was centrifuged to collect the “whole cell” fraction. The remaining cells were pelleted, resuspended in 50 mM Tris-HCl, 150 mM NaCl (pH 8), and lysed by sonication (5 min, 15s on/5s off, 40% amplitude). Cell debris was removed by centrifugation at 5000 xg for 15 min, and the supernatant was centrifuged at 175,000 xg for 45 min to separate membrane and soluble fractions. The membrane pellet was resuspended in 500 µL Tris buffer with 0.05% dodecylmaltoside.

Proteins in the supernatant were precipitated using 40% trichloroacetic acid, incubated on ice for 15 min, and pelleted by centrifugation at maximum speed for 20 min. The pellet was resuspended in 90 µL SDS loading dye and neutralized with 10 µL of saturated Tris solution. Proteins from the supernatant, membrane, and whole cell fractions were separated on two 15% SDS-polyacrylamide gels. One gel was stained with Coomassie blue, and the other was used for Western blotting with anti-HA antibodies, as described for the cleavage assays.

### Electronic Microscopy

Cells were stressed as described above. Five μL of culture were spotted onto a copper-coated grid and incubated for 30 seconds. The liquid was removed using Whatman paper, and this was repeated five times. The grid was then washed with 20 μL of MilliQ water and dried with Whatman paper. A drop of 2% uranyl acetate was added to the grid and immediately removed with Whatman paper, repeating the staining step twice. After removing excess liquid, the grid was air-dried. Imaging was performed using a Hitachi HT7800 operated at 100 kV, equipped with an EMSIS Xarosa 20-megapixel CMOS camera.

### DNA exchange Assays

*S. acidocaldarius* mutants were assayed for DNA exchange using MW001 (Δ91-421 nt *pyrE*) and JDS22 (Δ16-38 nt *pyrE*) and derived mutants. For DNA exchange, cells were grown to an OD of 0.4-0.5, spun down at 75°C for 20 minutes at 3,000 × g, and resuspended in warm Brock media without uracil to a theoretical OD of 0.7. Ten mL of culture were stressed with UV light (75 J/m^2^). Cultures were mixed (1 mL of each mating partner) and incubated for 4 hours at 75°C in a 24-well plate. Prototrophic recombinants (*pyr*+) were selected by plating the mixes on plates without uracil and incubating for 5-6 days at 75°C. The OD600 of mixed cultures was measured before plating for standardization. Colony PCR was performed on the resulting colonies using primers listed in Table S2.

## Conflict of interest

The authors declare that the research was conducted in the absence of any commercial or financial relationships that could be construed as a potential conflict of interest.

### Authors contributions

AR, AW and MvW performed bioinformatics analysis. AR, AW, CK, BW and NF obtained the deletion mutants. AR, AW, CK, NF and RT performed cell aggregation assays. AS constructed expression plasmids and performed cleavage assays. AR did grid preparation for EM and SS performed EM imaging. AR, AW, TNL, NF, RT, BW and MvW performed DNA exchange experiments. AR supervised NF, RT and AS and performed protein localization. AR and MvW prepared the figures and wrote the manuscript. AR, AW, MvW and SVA designed experiments and edited the manuscript.

## Acknowledgments

We would like to thank the EM facility at the Faculty of Biology, University of Freiburg, for access to the TEM for generation of data. The TEM (Hitachi HT7800) was funded by the DFG grant (project number 426849454) and is operated by the University of Freiburg, Faculty of Biology, as a partner unit within the Microscopy and Image Analysis Platform (MIAP) and the Life Imaging Center (LIC), Freiburg.

AR received support from the VW Momentum grant 94993. AW was funded by the DFG Grant AL1206. SS and SVA were supported by the Collaborative Research Centre SFB1381 funded by the Deutsche Forschungsgemeinschaft (DFG, German Research Foundation), Project-ID 403222702—SFB 1381 and by the Deutsche Forschungsgemeinschaft (DFG, German Research Foundation) under Germany’s Excellence Strategy (CIBSS – EXC-2189 – Project ID 390939984). TNL was supported by a 911 grant from the Ministry of Education and Training, Vietnam. MvW was funded by VW Momentum grant 94993.

## Notes

### Competing Interest Statement

The authors have declared no competing interest.

