## Supplementary material for "New components of the community based DNA-repair mechanism in Sulfolobales"

**Table S1.** Homologs of proteins of the *S. acidocaldarius* iModulon found in other Thermoproteota

| | Ced system | | | | | Ups | | | | | | | |
|---|---|---|---|---|---|---|---|---|---|---|---|---|---|
| **Organism** | **CedA1** | **CedA** | **CedA2** | **CedB** | **CedD** | **UpsA/B** | **UpsC** | **UpsX** | | | | | **TFB3** |
| **Sulfolobales** | | | | | | | | | | | | | |
| *Sulfolobus acidocaldarius* DSM639 | Saci_0569 | Saci_0568 | Saci_0567 | Saci_0748 (TMD) | **Saci_0667** (TMD) | **Saci_1496a/b** | **Saci_1302** | Saci_1493 | **Saci_1270** | **Saci_0951** | **SACI_RS06010** | **Saci_1225** | **Saci_0665** |
| *Saccharolobus islandicus* REY15A | SiRe_1316 | SiRe_1317, SiRe_2100 | SiRe_1318 | SiRe_1857 (TMD) | SiRe_1715 (TMD) | SiRe_1881 | SiRe_1957 | SiRE_1878 | SiRe_0137 | SiRe_0187 | SiRe_0269 | -- | SiRe_1717 |
| *Saccharolobus solfataricus* P2 | + | SSO0691, SSO3146 | + | SSO0152 (TMD) | SSO0283 (TMD) | SSO0117-8 | SSO0037 | SSO0121 | SSO2338 | SSO2395 | + Sso2478-79 | -- | SSO0280 |
| *Sulfurisphaeras tokodaii* str. 7 | + | ST0407 | STS057 | ST0197 (TMD) | ST0320 (TMD) | ST1399 | STK_16160 | ST1396 | STK_25810 | STK_0528 | + Before STK_0968 | STK_0767 | STK_0322 |
| *Acidianus hospitalis* W1 | Ahos_0872 | Ahos_0873 | + | Ahos_0881 (TMD) | Ahos_0795 (TMD) | -- | | AHOS_0568 | Ahos_0115 | AHOS_2190 | AHOS_1478 | -- | AHOS_0797 |
| *Metallosphaera cuprina* Ar-4 | Mcup_1940 | Mcup_1939 | Mcup_1938 | Mcup_1911 | Mcup_2010 | Mcup_0181/0 | Mcup_0609 | Mcup_0184 | | | Mcup_0645 | -- | Mcup_2012 |
| *Metallosphaera sedula* DSM 5348 | Msed_0119 | Msed_0120 | + | Msed_0162 (1TMD) | Msed_2263 (1TMD) | Msed_2106/7 | Msed_1620 | Msed_2103 | Msed_0293 | | Msed_1587 | -- | Msed_2265 |
| **Desulfurococcales** | | | | | | | | | | | | | |
| *Aeropyrum pernix* | APE_0220a | APE_0220 | - | APE_0219 | APE_2093 | | | | | | | | |
| *Aeropyrum camini* | ACAM_0169 | ACAM_0168 | - | ACAM_0167 | ACAM_1311 (TMD) | | | | | | | | |
| *Hyperthermus butylicus* DSM 5456 | + | Hbut_1291 | + | Hbut_1292 | Hbut_0386 | | | | | | | | Hbut_1563 |
| *Ignicoccus hospitalis* KIN4/I | Igni_0950 | Igni_0716 | - | Igni_0715 | Igni_0221 | | | | | | | | |
| *Ignisphaera aggregans* DSM 17230 | Igag_0133 | Igag_0132 | + | Igag_0989 | Igag_0110 | | | | | | | | Igag_0032 |
| *Pyrolobus fumarii* 1A | Pyrfu_1644 | Pyrfu_1645 | Pyrfu_1646 | Pyrfu_1647 | Pyrfu_0100 (TMD) | | | | | | | | |
| *Staphylothermus hellenicus* DSM 12710 | Shell_1453 | Shell_1452 | Shell_1451 | Shell_1450 | Shell_1558 | | | | | | | | Shell_1532 |
| *Staphylothermus marinus* F1 | Smar_1011 | Smar_1012 | + | Smar_1013 | Smar_0908 | | | | | | | | Smar_0933 |
| *Thermogladius cellulolyticus* 1633 | TCELL_0318 | TCELL_0319 | TCELL_0320 | TCELL_0321 | TCELL_0285 | | | | | | | | |
| *Thermosphaera aggregans*, DSM 11486 | Tagg_0840 | Tagg_0841 | Tagg_0842 | Tagg_0843 | Tagg_1287 | | | | | | | | Tagg_1257 |
| **Acidilobales** | | | | | | | | | | | | | |
| *Acidilobus saccharovorans* 345-15 | ASAC_1270 | ASAC_1271 | - | ASAC_1272 | Asac_0325 | | | | | | | | |
| | Ted system | | | | | | | | | | | | |
| | **TedC** | **TedA** | | **TedB** | **TedD** | | | | | | | | |
| **Thermoproteales** | | | | | | | | | | | | | |
| *Caldivirga maquilingensis* | Cmaq_1736 | Cmaq_1575 | | Cmaq_0645 (TMD) | Cmaq_1723 | | | | | | | | |
| *Pyrobaculum calidifontis* | Pcal_0765 | Pcal_0766 | | Pcal_0756 (TMD) | Pcal_0763 | | | | | | | | |
| *Pyrobaculum oguniense* | Pogu_0631 | Pogu_0632 | | Pogu_0536 | Pogu_0622 | | | | | | | | |
| *Vulcanisaeta distributa* | Vdis_0149 | Vdis_0150 | | Vdis_0137 | Vdis_0134 | | | | | | | | |
| *Vulcanisaeta moutnovskia* | Vmut_1011 | Vmut_1012 | | Vmut_1001 (TMD) | VMUT_0996 | | | | | | | | |

+: Homologous protein found but not annotated
-: No homologous protein found
TDM: N-terminal transmembrane domain
Underlined are genes next to each other on the genome: operon?

**Table S2.** Plasmids and primers used in this study

| **Plasmid** | ***Backbone*** | ***Use*** | ***Reference*** |
|---|---|---|---|
| pSVA431 | | Backbone for deletion plasmids | (Wagner et al., 2012) |
| pSVAaraFX-Stop | | Backbone for expression plasmids | (van der Kolk et al., 2020) |
| pSVAaraFX-HA | | Backbone for expression plasmids | (van der Kolk et al., 2020) |
| pSVA914 | | Base vector for cleavage assay. Contains PibD under IPTG promoter | (Henche et al., 2014) |
| pSVA3507 | pSVA431 | Plasmid for in-frame deletion of *saci_0667* (*cedD*) | This work |
| pSVA12831 | pSVA431 | Plasmid for in-frame deletion of *saci_1302* (*upsC*) | This work |
| pSVA6579 | pSVA431 | Plasmid for in-frame deletion of *saci_0951* | This work |
| pSVA12830 | pSVA431 | Plasmid for in-frame deletion of *saci_1225* | This work |
| pSVA12829 | pSVA431 | Plasmid for in-frame deletion of *saci_1270* | This work |
| pSVA12832 | pSVA431 | Plasmid for in-frame deletion of *saci_RS06010* | This work |
| pSVA13161 | pSVAaraFX-HA | Expression of *saci_1302* (*upsC*), HA-tagged | This work |
| pSVA13168 | pSVAaraFX-Stop | Expression of *saci_1302* (*upsC*) | This work |
| pSVA13169 | pSVA914 | For cleavage assay of truncated *saci_1302* in *E. coli* | This work |
| pSVA13197 | pSVA13169 | Plasmid derived from pSVA13169 without *pib*D | This work |

| **Primer No.** | **Sequence (5´ to 3´)** | **Use** |
|---|---|---|
| 12402 | CATGCTCGAGGAATTGGCTGAGTCAGTTAC | US *saci_1302* (*upsC*) fw *XhoI* |
| 12403 | GAATAAAAATAATGTCGTTTGAATTCTTGTTTTAATAAACAGATG | US *saci_1302* (*upsC*) rv overlap |
| 12404 | CAAGAATTCAAACGACATTATTTTTATTCTTATTTTTCAC | DS *saci_1302* (*upsC*) fw overlap |
| 12405 | CATGGGCCCTCGCAAAGCTGATTTATTCTTACC | DS *saci_1302* (*upsC*) rv *ApaI* |
| 12076 | CTGGAAAACACGCCGTTAAAGAT | Fw sequencing of *saci_1302* (*upsC*) deletion |
| 12077 | CTGGTTAAGAATGCCTTCTCGATC | Rv sequencing of *saci_1302* (*upsC*) deletion |

| | | |
|---|---|---|
| 12086 | ATATATGCTCTTCGAGTAAAAGGAGAAAGAAGGGAATATCGACA | Cloning of *saci_1302* (*upsC*) Fw |
| 12087 | TATATAGCTCTTCATGCACCTGTTGAGACGTCGATAATATAGTC | Cloning of *saci_1302* (*upsC*) Rv |
| 13221 | ATATATGCTCTTCTAGTAAAAAGGGAATATCATCAATCCTAGGT | Cloning of *SSO0037* ($upsC_{Sa.sol}$) Fw |
| 13222 | TATATAGCTCTTCATGCCACAGCAAAGTAAGGATAGTAGACATC | Cloning of *SSO0037* ($upsC_{Sa.sol}$) Rv |
| 13223 | ATATATGCTCTTCTAGTAGAGGTATATCTAGTATTTTAGGTACC | Cloning of *ST1616* ($upsC_{S.tok}$) Fw |
| 13224 | TATATAGCTCTTCATGCACTCTGCTGGACTTGTAATAGATATAC | Cloning of *ST1616* ($upsC_{S.tok}$) Rv |
| 13225 | ATATATGCTCTTCTAGTAAAAAGGGAATATCGTCAATCTTAGGT | Cloning of *SiRe_1957* ($upsC_{S.isla}$) Fw |
| 13226 | TATATAGCTCTTCATGCAGGATAATAGACGTTCTCAGGGATCAT | Cloning of *SiRe_1957* ($upsC_{S.isla}$) Rv |
| 12094 | GGAGGAATTAACCATGAAGAGAAGGAAGAAGGGAAT | Fw gene truncated *saci_1302* (*upsC*) for *in vivo* assembly |
| 12095 | GTAGGATCCCCCGGGGGAGATTAAAGAAGATGGTG | Rv gene truncated *saci_1302* (*upsC*) for *in vivo* assembly |
| 12096 | CTTTAATCTCCCCCGGGGGATCCTACCCGTATGACGT | Fw plasmid pSVA914 for *in vivo* assembly |
| 12097 | CTTCTCTTCATGGTTAATTCCTCCTGTTAGCCCAA | Rv plasmid pSVA914 for *in vivo* assembly |
| 13235 | GGCATGCATAATGTGCCTGTCAAATG | Fw for pSVA13197 to delete *pibD* |
| 13236 | TGCATGCCGCTTCGCCTTCG | Rv for pSVA13197 to delete *pibD* |
| 5610 | GCGCATATGAGTTGGATAAGGAAGAAAAG | US *saci_0667* fw *NdeI* |
| 5611 | GAAGAAGACTAAGCTAAAGAGTAACTACTTATTTATGAG | US *saci_0667* (*cedD*) rv overlap |
| 5612 | TTACTCTTTAGCTTAGTCTTCTTCTTTACCTTTTTC | DS *saci_0667* (*cedD*) fw overlap |
| 5613 | GCGGGATCCAACAGTTGCAGATGTAGTTAG | *DS saci_0667* (*cedD*) rv *BamHI* |
| 5614 | GGCGACAAGGTTAGGAAGAGACG | Fw sequencing of *saci_0667* (*cedD*) deletion |
| 5615 | GAGCATTAAGAGATGCCTTAG | Rv sequencing of *saci_0667* (*cedD*) deletion |
| 9494 | CATGCTCGAGGTGGCTTTCTTACCTTCAAC | US *saci_1270* fw *XhoI* |
| 9495 | GATTAATTGAGTTTATTGTACTTACTTAGGTCATTAAC | US *saci_1270* rv overlap |
| 9496 | TGACCTAAGTAAGTACAATAAACTCAATTAATCAC | DS *saci_1270* fw overlap |
| 9497 | GATCGGGCCCATGGACATGAGTACCTTAAC | DS *saci_1270* rv *ApaI* |
| 12080 | TCAATCAACTCTGCTCTAAC | Fw sequencing of *saci_1270* deletion |
| 12081 | CTTTACTCGCCAATAAATCC | Rv sequencing of *saci_1270* deletion |
| 9498 | CATGCTCGAGAATAAGCAAGTCCCTAAGTG | US *saci_1225* fw *XhoI* |

| | | |
|---|---|---|
| 9499 | GTAATAAACACGTTTAGGTCTATTTCAGCATTTTTAAATGTC | US *saci_1225* rv overlap |
| 12400 | AAATGCTGAAATAGACCTAAACGTGTTTATTACTG | DS *saci_1225* fw overlap |
| 12401 | GCATGGGCCCGGAAGCACTTAGGATAATTG | DS *saci_1225* rv *ApaI* |
| 12082 | GGAGACAGTACTTCAAATTC | Fw sequencing of *saci_1225* deletion |
| 12083 | CTGCACAATCCACACTAATG | Rv sequencing of *saci_1225* deletion |
| 12406 | CATCTCGAGTTACTACAGAAGGCAATTTAACTCAAG | US *saci_RS06010* fw XhoI |
| 12407 | AGTCTTCAAGAACTTATAAATCCTTTGTTCTAAAAATATC | US *saci_RS06010* rv overlap |
| 12408 | CTGATATTTTTAGAACAAAGGATTTATAAGTTCTTGAAGACTTCAG | DS *saci_RS06010* fw overlap |
| 12409 | CATGGGCCCGGTATCAAAGAACCAAGAAAG | DS *saci_RS06010* rv *ApaI* |
| 12078 | AGAGTCACATGCTACCTATC | Fw sequencing of *saci_RS06010* deletion |
| 12079 | AGCCCAAACCTTTAGTTGAG | Rv sequencing of *saci_RS06010* deletion |
| 12410 | CATCTCGAGCTCCCACGTCCTAAAGCCATAACCAAG | US *saci_0951* fw XhoI |
| 13976 | CATCATGTCCTCGTGTCATCCATATTACTCTTATTGAGTT | US *saci_0951* rv overlap |
| 13977 | GAGTAATATGGATGACACGAGGACATGATGTATAAATTAA | DS *saci_0951* fw overlap |
| 12413 | CATGGGCCCGAAATTCCTCAATGTATGTTAATGCGTTCTC | DS *saci_0951* rv *ApaI* |
| 12084 | CCTGCAACTACCCAATATTC | Fw sequencing of *saci_0951* deletion |
| 12085 | TCCTAAATGAGTTCGGTATG | Rv sequencing of *saci_0951* deletion |
| 13946 | ATCGGGGCCCAAAGCTTTCAATTAAATACGTATC | Fw US *saci_0569* (*cedA1*) ApaI |
| 13947 | AAAAGGATTTAGCATTTATCATTCAGGATAAATATTTAA | Rv US *saci_0569* (*cedA1*) overlap |
| 13948 | TATCCTGAATGATAAATGCTAAATCCTTTTCAACTTTT | Fw DS *saci_0569* (*cedA1*) overlap |
| 13949 | ATGCCCATGGGGTAAACCTACATAAAAGACTAT | Rv DS *saci_0569* (*cedA1*) NcoI |
| 13950 | ATCGGGGCCCGTTAAAACAATATTAACTGCCAG | Fw US *saci_0567* (*cedA2*) ApaI |
| 13951 | CTAAGAAAAAAGCTTTGTTCTCACCGTTTCCTAAAATAC | Rv US *saci_0567* (*cedA2*) overlap |
| 13952 | GAAACGGTGAGAACAAAGCTTTTTTCTTAGCTTCTTTC | Fw DS *saci_0567* (*cedA2*) overlap |
| 13953 | ATGCCCATGGCTTAAAATATAAGTTCCATCGTT | Rv DS *saci_0567* (*cedA2*) NcoI |
| 2010 | GTAGGGCCCGTGTATAATGATGACCTATTTAGCTG | Amplification of *upsE* Fw |
| 2015 | GTAAACTGGAAGCCTATAAGG | Amplification of *upsE* Rv |
| 5608 | CGGTAAGATTCTTGTAATGGGCGCAGGTAG | Amplification of *cedB* Fw |

| | | |
|---|---|---|
| 5609 | CTACTTTGTTATGCATAGCTAATAGCC | Amplification of *cedB* Rv |
| 6902 | CTTTGCGGAGAGGTATTCAG | qPCR *herA* (*saci_0053*) Fw |
| 6903 | GCTACTATGGCGTCTATTGC | qPCR *herA* (*saci_0053*) Rv |
| 13268 | AACCCGTTTCTCTGAAGG | qPCR *nurA* (*saci_0050*) Fw |
| 13269 | AGGCCACCAACTTATGTC | qPCR *nurA* (*saci_0050*) Rv |
| 1480 | CCTGCAACATCTATCCATAACATACCGA | qPCR *secY* fw |
| 1481 | CCTCATAGTGTATATGCTTTAGTAGTAG | qPCR *secY* rv |
| 2075 | GCTAGTAAAGCCAACAAGAGTG | qPCR *upsE* fw |
| 2076 | ATATAGTCGCTGCTACCCTATG | qPCR *upsE* rv |
| 3034 | GGAGGAGTGTTCAGTTCTTG | qPCR *cedA* fw |
| 3035 | GGTGGTGCCTTTATAGGAAGTG | qPCR *cedA* rv |
| 2069 | GCTCTACTTGATAAAAAATTGC | *pyrE* check fw |
| 2070 | GCTCTGATGTATCCCATAGG | *pyrE* check rv |

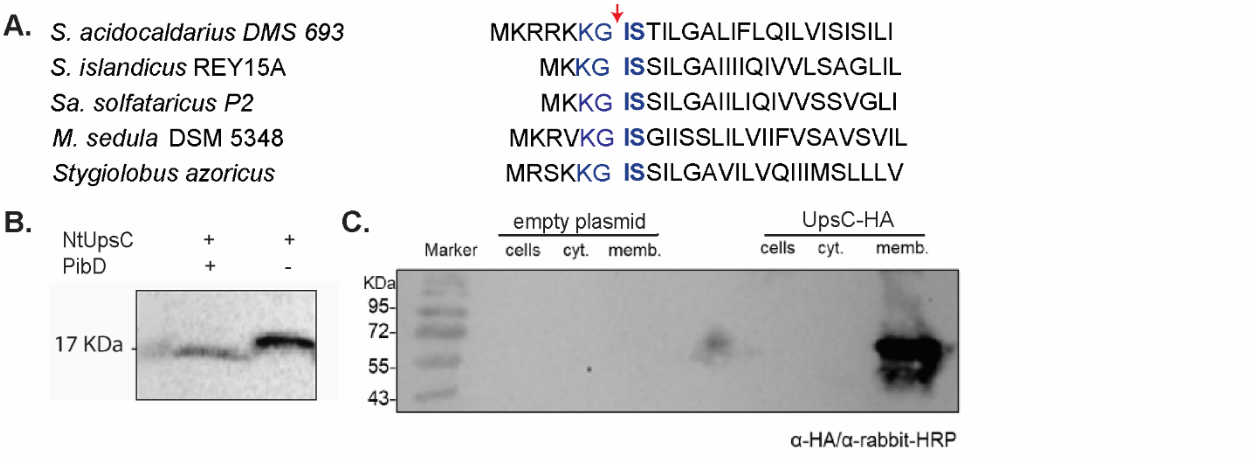


**Figure S1:** **(A)** Alignment of the N-terminus of UpsC from different Sulfolobales. **(B)** Cleavage assay via heterologous expression of the N-terminal portion of UpsC and PibD in *E. coli*. **(C)** Localization of a tagged version of UpsC in the membrane fraction of cells.

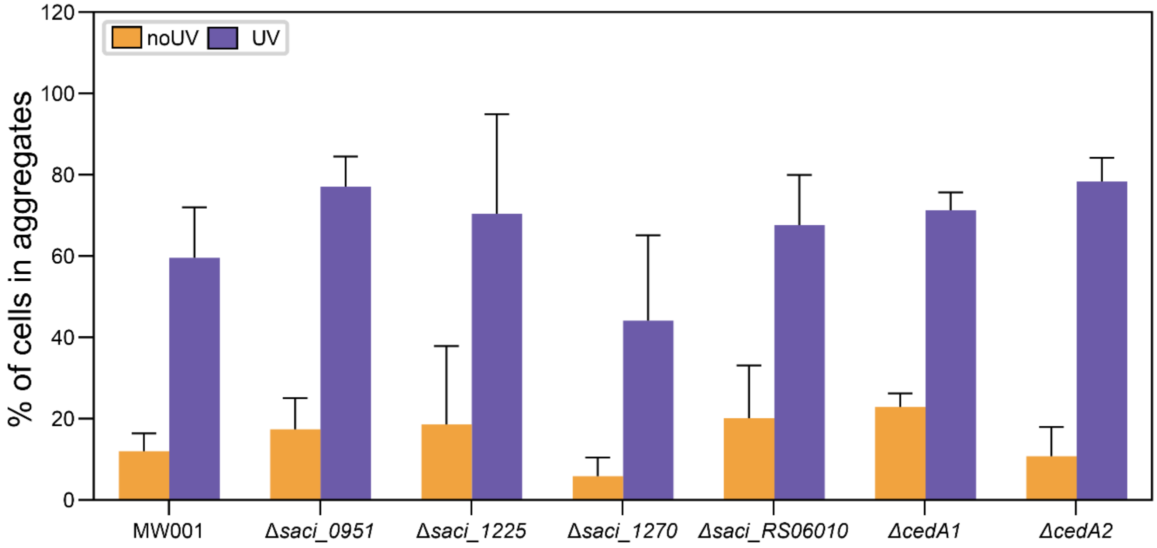


**Figure S2:** Aggregation assays on *S. acidocaldarius* MW001, *Δsaci_0951, Δsaci_1225, Δsaci_RS06010, ΔcedA1 ΔcedA2.* Shown are percentages of aggregated cells, after UV light exposure and the non-exposed control

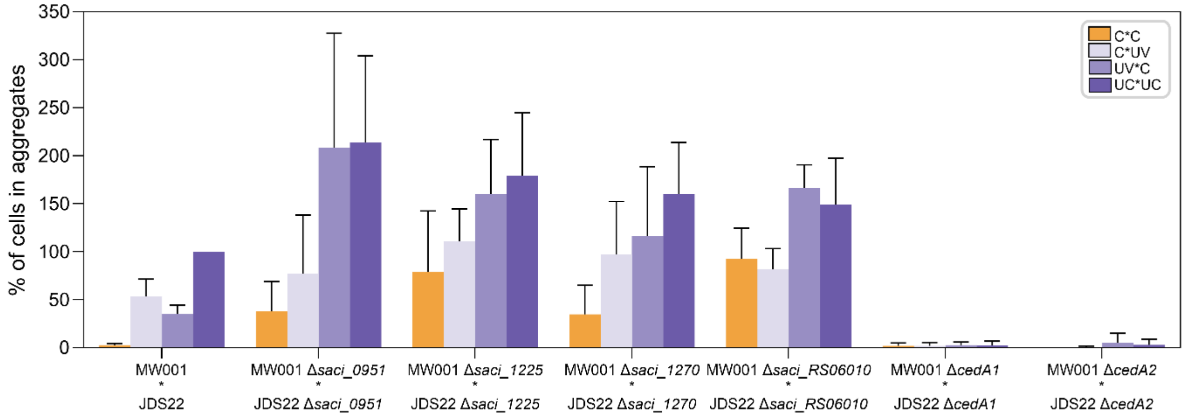


**Figure S3:** DNA exchange assays with *Δsaci_0951, Δsaci_1225*, Δ*saci_1270*, Δ*saci_RS06010*. Two different mutants (either in JDS22 or MW001 backgrounds), exposed to UV light (UV) or left untreated (C), were mixed in the indicated combinations and plated on selective media. Both background strains carried mutations in the *pyrE* gene, which is involved in uracil biosynthesis. The mutations were located at different genomic positions, allowing recombination between the strains to restore the wild-type *pyrE* phenotype. The bars represent the average of at least three independent mating experiments, with each experiment normalized to the JDS22 (UV) * MW001 (UV) condition, set at 100%.
